# Fetoplacental circadian rhythms develop and then synchronize to the mother *in utero*

**DOI:** 10.64898/2025.12.17.695051

**Authors:** K.L. Nikhil, Keenan Bates, Elizabeth Sapiro, Jacob L. Amme, Ronald McCarthy, Sarah L. Speck, Varun Vasireddy, Ethan Roberts, Carmel A. Martin-Fairey, Miguel-E. Domínguez-Romero, Sandra Paola Cárdenas-García, Sarah K. England, Erik D. Herzog

## Abstract

Circadian rhythms in gene expression and hormones are ubiquitous across species and differentiated cell types, yet their developmental origins remain poorly understood. This study aimed to determine if daily rhythms can be detected *in utero* and if they synchronize to the mother. We developed methods to longitudinally monitor PERIOD2 (PER2), a core circadian clock protein, from embryonic day (E)8.5 to E17.5 by restricting PER2::LUCIFERASE (PER2::LUC) expression to the mouse fetoplacental unit (fetus and fetal-derived tissues). *In utero* fetoplacental bioluminescence imaging showed that PER2 levels rose exponentially during pregnancy, with variable daily peak times that stabilized to dusk by E15.5. Interestingly, pregnancies that did not exhibit daily *in utero* PER2 variation were more likely to fail. Because maternal glucocorticoids have been implicated in fetal development and synchronizing other circadian tissues, we tested its capability to shift fetoplacental PER2 rhythms *in utero*. Daily subcutaneous glucocorticoid injections over five days of late pregnancy phase-dependently shifted the fetoplacental PER2 rhythms *in utero*. Blocking glucocorticoid signaling *in vitro* reduced synchrony between maternal and fetal placenta by ∼40%. We conclude that *in utero* daily rhythms gradually develop and synchronize with the mother prior to birth, potentially through glucocorticoid signaling.

## Introduction

Circadian rhythms in gene expression and hormones generate daily cycles in behavior and physiology in many species and cell types (R. Zhang et al., 2014). In mammals, individual cells utilize transcription-translation feedback loops of core clock genes, including *Period2* (*Per2)*, to generate and maintain circadian rhythms (Cox & Takahashi, 2019). In the adult, the suprachiasmatic nucleus (SCN) in the hypothalamus generates near-24 h rhythms, entrains to environmental cues such as the daily light-dark cycle, and regulates daily rhythms throughout the body (Herzog, 2007). When and where these daily molecular oscillations first arise in development is debated.

Daily rhythms in clock genes are not found in embryonic stem cells or induced pluripotent stem cells until differentiation (Kowalska et al., 2010; Umemura et al., 2017a; Yagita et al., 2010). Dedifferentiation back into the pluripotent stem cell state causes these rhythms to be lost (Yagita et al., 2010). *Ex vivo* studies have found circadian rhythms in a variety of fetal tissues (Carmona-Alcocer et al., 2018; Dolatshad et al., 2010a; Landgraf, Achten, et al., 2015; Wreschnig et al., 2014). For example, explants of fetal mouse liver and kidneys exhibit daily rhythms as early as embryonic day (E)13.5 (mouse gestation lasts 19-20 days) when the SCN does not yet exhibit rhythms. A day later, ∼10% of cells within the fetal mouse SCN develop circadian rhythms and, by E15.5, almost all SCN cells show synchronized daily rhythms (Carmona-Alcocer et al., 2018; Landgraf, Achten, et al., 2015; Wreschnig et al., 2014). However, *in vivo* data are not fully concordant. Early *in utero* studies of fetal rat SCN showed day-night differences in metabolic activity from E19 (rat gestation lasts 21-22 days) (Reppert & Schwartz, 1983, 1984). *In vivo* or *in vitro* measurements of mRNA or protein expression of clock genes in the fetal rat SCN, liver or whole mouse embryos either failed to detect rhythms or only detected rhythms one or two days prior to birth (Dolatshad et al., 2010a; Houdek & Sumová, 2014; Kováciková et al., 2006; Sládek et al., 2004). This discrepancy may reflect differences between *in vitro* and *in vivo* induction of circadian cycling. However, prior *in vivo* approaches required sacrificing many animals and pooling data from multiple fetuses and pregnancies. This method can detect when fetal rhythms are synchronized across pregnancies but cannot assess when individual pregnancies develop rhythms. One prior study assessed fetal clock gene expression in rats using a real time reporter of clock gene Period1 transcription, but did not have the temporal resolution to determine if or when daily rhythms develop (Saxena et al., 2007). Due to these inconsistent findings between studies, when circadian rhythms initiate in the fetus remains unknown.

Pregnancy offers an interesting scenario as maternal and fetal circadian oscillators likely interact as fetal circadian rhythms develop. Previous research has indicated that maternal signals can drive rhythms in the fetal SCN (Greiner et al., 2022). Maternal humoral signals, such as melatonin or glucocorticoids, may act to entrain or drive daily rhythms in the fetus (Čečmanová et al., 2019; Davis & Mannion, 1988). Any humoral signal that synchronizes offspring *in utero* to daily environmental cues must pass through the placenta to interact with the fetus. Importantly, *in vivo* and *in vitro* experiments demonstrate that the placenta also exhibits circadian rhythms in clock genes and enzymes (Akiyama et al., 2010; Čečmanová et al., 2019; Crew et al., 2018; Wharfe et al., 2011). For example, daily rhythms in placental 11β-hydroxysteroid dehydrogenase (11β-HSD2) inactivate glucocorticoids during rest, allowing active glucocorticoids to cross the placenta during wake, thereby generating a signal that could entrain the fetus (Burton & Waddell, 1999; Čečmanová et al., 2019; Holmes et al., 2006). Importantly, some of the potential entraining signals function in other developmental processes, such as glucocorticoid regulation of fetal lung development (Bird et al., 2015). In pregnancies at risk for preterm birth, antenatal glucocorticoid treatments are used to accelerate fetal lung development (Bird et al., 2015; Briceño-Pérez et al., 2019; Roberts et al., 2017). However, the time-of-day of treatment is rarely considered, which could reduce efficacy or drive abnormal circadian rhythm development in the fetus. Studies in mice (daily injections from E11.5 onward) and humans (2 daily doses at 24-34 weeks of gestation) showed that glucocorticoid treatments during gestation given out-of-phase relative to the endogenous rhythm caused diminished resilience to stress for the offspring (Astiz et al., 2020a). Therefore, more research needs to be done to understand how the timing of glucocorticoid administration affects the development of fetal circadian rhythms.

This study aimed to measure maternal-fetal coordination of clock gene expression during pregnancy. We hypothesized that circadian rhythms arise in the fetoplacental unit (fetus and placenta) early in development and synchronize to the mother. By monitoring clock gene expression in pregnant females expressing PER2::LUCIFERASE (PER2::LUC) exclusively in the fetoplacental tissue, we observed the gradual emergence of synchronized daily rhythms in PER2 that stabilized around E15.5. Independent *in vitro* recordings from maternal tissues expressing PER2::LUC revealed parallel changes in daily PER2 peak times in the uterus and the placenta. While both maternal and fetal layers of the placenta exhibited intrinsic circadian rhythms beginning as early as E9, daily glucocorticoid administration to dams increased synchrony across the maternal-fetal tissues while glucocorticoid receptor antagonists, *in vitro,* disrupted synchrony within the placenta. These results support the hypothesis that fetal circadian rhythms develop during pregnancy and entrain to the mother prior to birth, potentially through glucocorticoid signaling.

## Results

### Successful pregnancies associated with development of high amplitude circadian rhythms in fetoplacental PER2

To characterize the development of fetal-derived clock gene expression *in utero*, we mated homozygous PER2::LUC males to wildtype female mice so that bioluminescence was derived exclusively from the fetoplacental units (Figure 1a, Supplementary Figure S1a). As a first step, we imaged ten anaesthetized dams following D-luciferin injections (150 mg/kg, 10 min prior imaging) every 12 h from ∼E8.5 to E14.5, and then every 4 h until E17.5 (Supplementary Figure S1b,c). We found that the four dams that successfully delivered exhibited an increase in the mean and day-night amplitude of *in utero* PER2::LUC bioluminescence. In contrast, six pregnancies did not show daily variations in PER2::LUC and resorbed the conceptus or failed to initiate labor (Supplementary Figure S1d-e).

**Figure 1.**
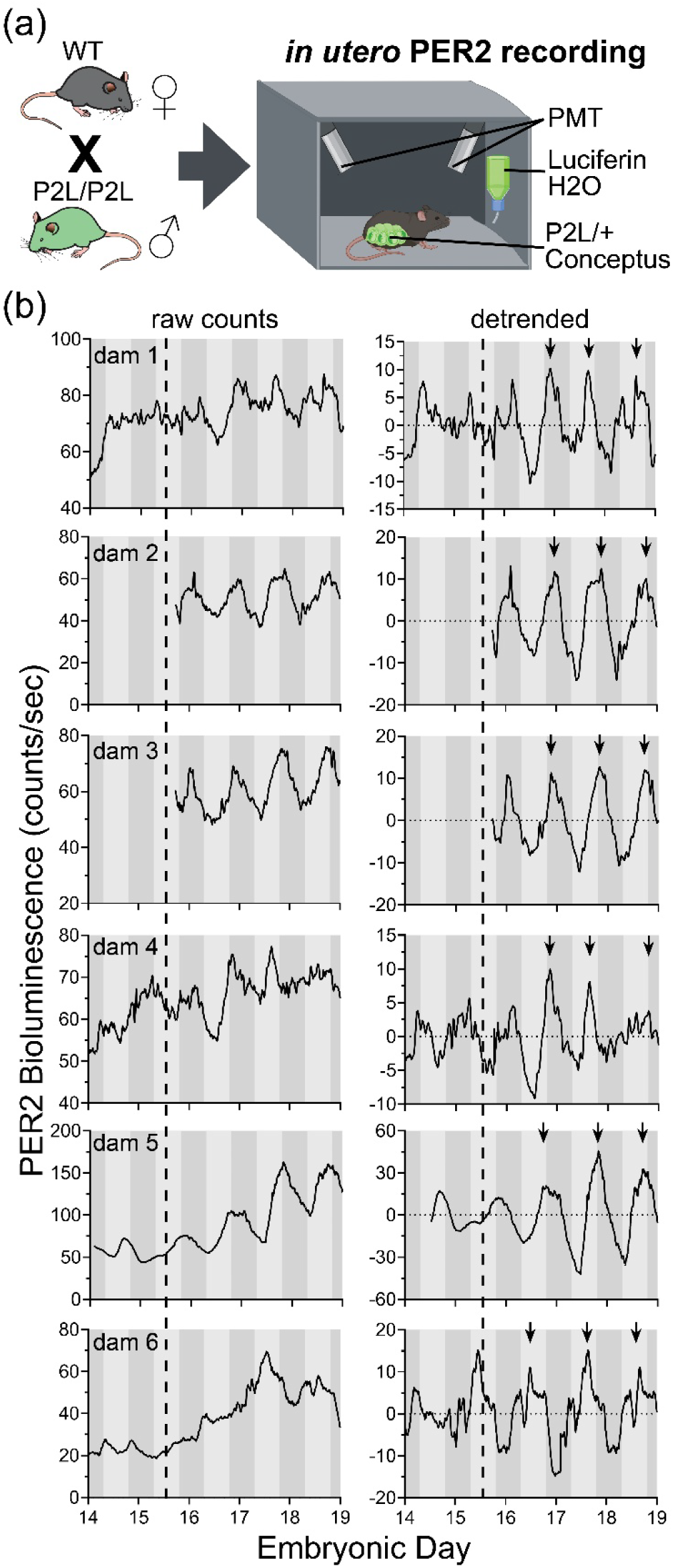
Daily fetoplacental PER2 bioluminescence gradually developed *in utero* and peaked near subjective dusk late in pregnancy. (a) Wild-type (WT) C57BL/6JN females were crossed with homozygous PER2::LUC (P2L) males to generate WT dams carrying heterozygous (P2L/+) fetuses, in which PER2-driven luciferase was expressed only in the fetoplacental units (fetuses and fetal-derived tissue). We recorded fetoplacental PER2 bioluminescence *in utero* from freely moving pregnant dams provided with 2.18 mM CycLuc1 in the drinking water in a light-tight constant dark chamber with two photomultiplier tubes (PMTs). (b) Raw counts (left) and detrended counts (right) of the fetoplacental PER2 bioluminescence recorded from all dams (n = 6) exhibited circadian rhythms embryonic day (E)15.5 onward (MetaCycle analysis, E15.5–E19.5), with daily PER2 peaks (arrows) occurring near subjective dusk during late gestation. Shading of the gray bars denotes the prior light and dark phases.

### Fetoplacental PER2 rhythms gradually developed and stabilized before birth

With a 60% pregnancy failure rate following repeated luciferin injections, we evaluated luciferin substrate delivery through drinking water as previously reported (Martin-Burgos et al., 2022; Sinturel et al., 2021). We longitudinally recorded bioluminescence every second in a freely moving homozygous PER2::LUC female mouse housed in a light-tight box with dual photomultiplier tubes. First, in a non-pregnant homozygous P2L female, we found that the daily PER2 peaked at dusk irrespective of whether luciferin (10 mM D-Luciferin in drinking water) was available only during daytime or *ad libitum* (Supplementary Fig. S2). Therefore, in a separate cohort, we provided wildtype females, pregnant with heterozygous PER2::LUC conceptus, continuous access to *ad lib* luciferin-supplemented water (2.18 mM CycLuc1) starting the day prior to recording (Figure 1a). Monitoring bioluminescence in four to six freely moving pregnant females from E14 to E19, we found that fetoplacental PER2 bioluminescence gradually increased through gestation, was circadian starting around E14.5-15.5 (Metacycle, *p* < 0.05) and reliably peaked around subjective dusk late in pregnancy (Figure 1b; *n* = 6 dams with 4-9 pups each).

In another cohort, we provided pregnant dams 10mM D-luciferin in drinking water (*ad lib*). We anaesthetized and imaged the dams every 12 h from ∼E8.5 to E14.5, and then every 4 h until E17.5 (Fig. 2a; Supplementary Fig. S3a). We again saw that *in utero* PER2 bioluminescence increased markedly through gestation and, by E15.5, showed significant circadian rhythms in 6 of the 7 pregnancies (*p* < 0.05, MetaCycle; Fig 2b, Supplementary Figure S3b). Again, the development of daily rhythms in fetal-derived tissues associated with successful delivery (Supplementary Tables S1 and S2). Using Rayleigh statistic (see Methods), we found that, starting around E15.5, the fetoplacental PER2 bioluminescence peaked at similar time of the day across independent dams (i.e., inter-pregnancy synchrony; Fig. 2d). Together, these findings indicate that fetoplacental PER2 circadian rhythms emerge early in pregnancy and progressively synchronize with maternal rhythms *in utero* to peak in the early night prior to birth.

**Figure 2.**
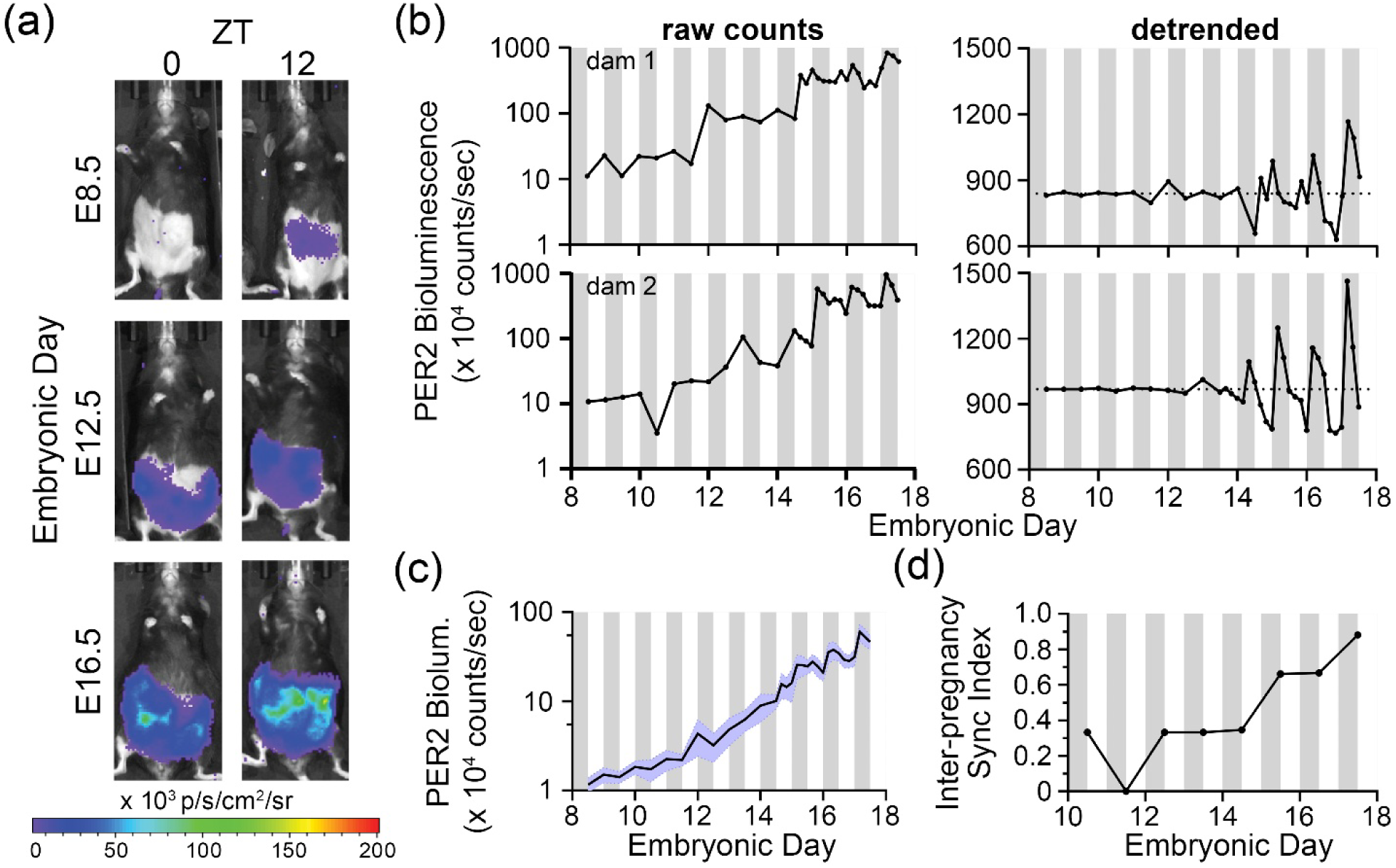
Fetoplacental PER2::LUC expression emerges early and grows in circadian amplitude across gestation. (a) Bioluminescence images captured through gestation at Zeitgeber Times (ZT) 0 (7:00am, lights on) and ZT12 (7:00pm, lights off) from a representative dam show that fetoplacental PER2 bioluminescence was detectable on embryonic day (E)8.5, increased through gestation, and was higher at ZT12 than at ZT0. (b) Raw (left) and detrended (right) fetoplacental PER2 bioluminescence imaged from two representative dams showing almost 100-fold increase in bioluminescence through gestation with daily peaks occurring reliably during the early night in the last days of gestation. White and gray bars indicate day and night. See Fig S3 for *in utero* imaging results of all dams. (c) Average fetoplacental PER2 bioluminescence showed daily rhythms starting around E15 when we initiated imaging every 4 hours. Error bars (blue) indicate SEM (n = 6 circadian dams). We excluded data from the one dam because her *in utero* bioluminescence did not score as circadian. (d) Inter-pregnancy sync index showed that the fetoplacental PER2 bioluminescence peaked at similar times across multiple dams as gestation advanced (n = 6 circadian dams). We quantified synchrony as the Rayleigh statistic computed from daily PER2 peak times across pregnancies and found a gradual increase in synchrony across dams in last stages of pregnancy. A sync index of 1 on a given day indicates fetoplacental PER2 bioluminescence peaked at the same time of day across all dams. White and gray bars indicate day and night, respectively.

### Glucocorticoid signaling synchronizes fetoplacental circadian rhythms

Past research has shown that glucocorticoids can act as an entraining signal in the fetal SCN (Čečmanová et al., 2019). Because glucocorticoids typically peak around dusk (∼ ZT13) in the mother (Cheifetz, 1971), we hypothesized that artificially elevating corticosterone levels around dawn would shift the time of fetoplacental peak PER2::LUC. A recent study showed that daily ZT0 corticosterone injections from E11.5-E15.5 caused maternal and fetal serum corticosterone to be higher at ZT1 than ZT13 (Astiz et al., 2020a). Using similar procedures, we injected 50 mg/kg corticosterone each day for five days at ZT0 from E13.5-E17.5 in seven pregnant wildtype mice expressing fetoplacental PER2::LUC. Bioluminescence imaging of fetoplacental PER2 starting on E8.5 until E17.5 revealed that PER2 was circadian by E15.5 in 5 of 7 corticosterone-injected dams (*p* < 0.05, MetaCycle; Supplementary Figure S4a). We noticed that four dams received their first corticosterone injection near the peak of their daily fetoplacental PER2::LUC expression and three dams received it around their trough (Supplementary Figure S4a). We found that daily corticosterone treatments that started near the peak or trough of fetoplacental PER2 bioluminescence on E13.5 (first day of injection), resulted in PER2::LUC rhythms that peaked 6-12 hours apart on E17 (Supplementary Figure S4b, c). The average difference in the peak PER2 time from E13 to E17 (i.e., phase shift) indicated a nearly 6-hour delay for pregnancies initially treated near the daily peak of fetoplacental PER2::LUC (Supplementary Figure S4c). Importantly, daily corticosterone injections starting on E13.5 increased synchronization of *in utero* PER2 rhythms (inter-pregnancy synchrony) earlier compared to control pregnancies (Supplementary Figure S4d vs Figure 2d).

We next evaluated whether corticosterone can synchronize PER2 rhythms across the fetoplacental tissue within each pregnancy (intra-pregnancy synchrony) by computing the Kuramoto order parameter based PER2 bioluminescence across the fetoplacental pixels (see Methods) as an indirect measure of intra-tissue synchrony. We first compared coherence in daily PER2::LUC expression measured across pixels and across 6 pup-sized regions-of-interest imaged within the uterus of representative pregnancies (Supplementary Figure S5). Both methods revealed similar intra-pregnancy increases in synchrony from E15 to 17. Pixel-based inter-pregnancy synchrony analysis of all pregnancies showed that, within each dam, those treated with ZT0 corticosterone (n = 5 dams with circadian *in utero* PER2::LUC, 406 ± 33 pixels/dam) increased synchrony across their fetoplacental pixels sooner than in controls (n = 5 vehicle-injected, 332 ± 103 pixels/dam, Figure 3, Supplementary Figure S6). These data indicate that morning corticosterone administration starting on E13 aligned peak fetoplacental PER2 rhythms across pups within a pregnancy within two days and accelerated circadian synchronization among fetuses.

**Figure 3.**
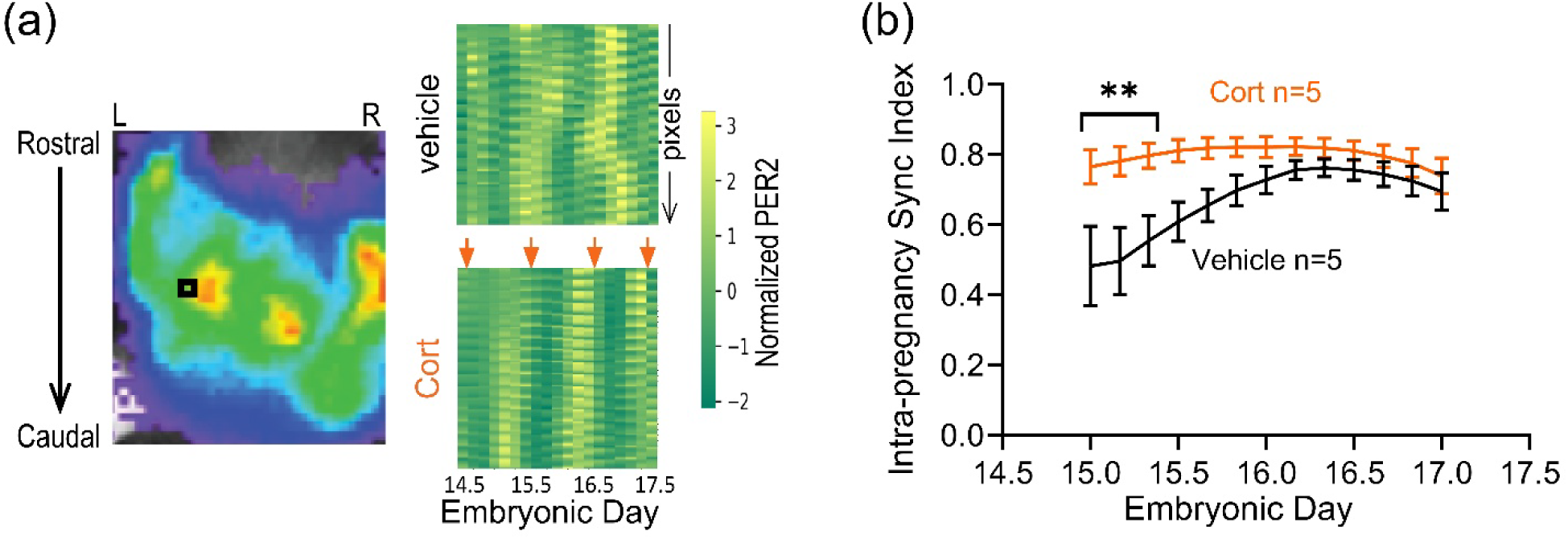
Corticosterone injections at dawn accelerated synchronization of *in utero* daily PER2::LUC rhythms within the fetoplacental regions. (a) PER2 bioluminescence imaged in the bilateral uterine horns of a representative dam (left) 10 min after injection of CycLuc1 (150 mg/kg body weight). The black square highlights one pixel over the left uterine horn adjacent to a bright fetus-sized area. Raster plots (right) show the normalized PER2 bioluminescence intensity of each pixel (row) recorded over three days within the uterus of representative dams injected daily at dawn (ZT0, arrows) with vehicle control (top) or corticosterone (Cort, bottom). Note that corticosterone injections increased the coherence of daily PER2 across pixels (intra-pregnancy sync). (b) Compared to the vehicle controls, daily corticosterone injections accelerated intra-pregnancy synchrony of PER2::LUC expression (synchrony among pixels within each pregnant uterus). We computed the sync index as the Kuramoto order parameter from the wavelet-based phase estimation of PER2 signals across fetoplacental pixels (332 ± 103 pixels/control and 406 ± 33 pixels/treated dam). Intra-pregnancy sync index (mean ± SEM) of control dams (black, n = 5 vehicle-injected) approached its maximum more gradually compared to that of corticosterone-treated dams (orange, n = 5; two-way ANOVA with Tukey’s multiple comparisons; \*\**p <* 0.01).

### Dynamic phase relationships between maternal and placental circadian tissues across development

We next tested for circadian coordination between maternal tissues and fetal placenta during pregnancy. From PER2::LUC homozygous females, we collected uterine, cervical, and placental explants at two estrous stages (metestrus and proestrus) and four embryonic ages (E9, E12, E15, and E18) and recorded their PER2::LUC expression *in vitro*. All tissues exhibited intrinsic circadian rhythms at each of the stages of pregnancy (Figure 4a). Although circadian periods did not dramatically change, daily amplitude of PER2 bioluminescence decreased with gestational age in the uterus (E9 vs E18, *p*=0.008) and placenta (E9 vs. E15, *p*=0.002 and E9 vs. E18, *p*=0.0005; Supplementary Figure S7a, b). In addition, PER2 bioluminescence in the uterus peaked around mid-morning during metestrus and mid-afternoon during proestrus (Figure 4b, Supplementary Figure S8), similar to the findings from prior research (Yaw et al., 2021). The cervix showed a similar shift in peak PER2 time from metestrus to proestrus (Figure 4b, Supplementary Figure S8). During pregnancy, the uterus consistently peaked about 2.5 h before the cervix from the same dam and between dams while its phase-relationship with the placenta varied through gestation (Figure 4b, c, Supplementary Figure S8). These results suggest that maternal tissues and fetal placenta possess intrinsic circadian rhythms, and their circadian amplitudes and phase relationships change over the course of healthy gestation.

**Figure 4.**
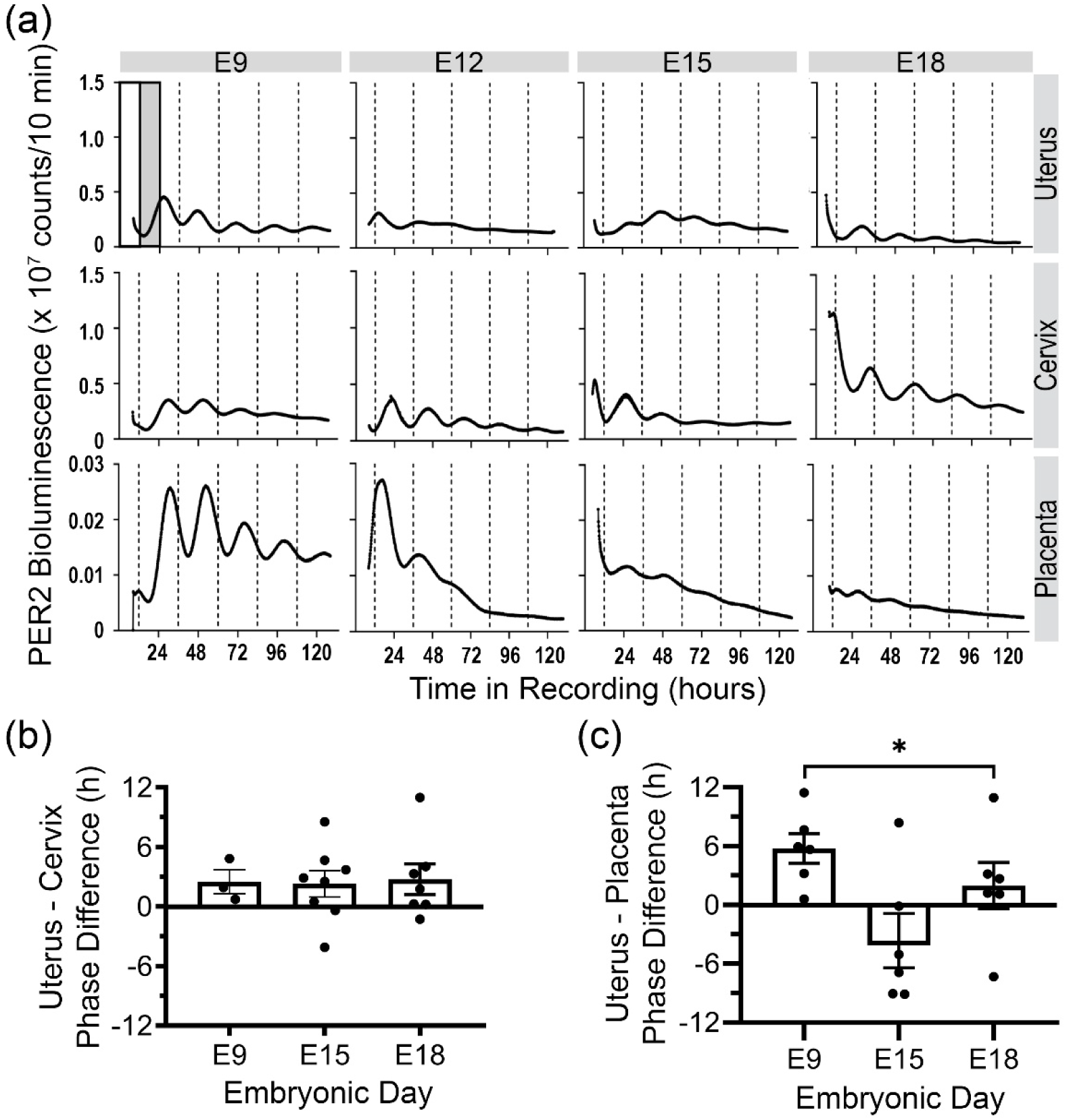
Coordinated daily rhythms intrinsic to the uterus, cervix, and placenta. (a) PER2 bioluminescence traces from representative uterus, cervix, and placental tissues exhibited circadian rhythms over 5 days of recording when explanted on embryonic days (E)9, E12, E15, or E18. Relative to the prior light–dark cycle (white and gray bars), the daily PER2 bioluminescence peak times depended on both tissue type and gestational stage. (b) Within the same dam, the uterus peaked ∼2.5 h before the cervix across all stages. (c) The phase relationship between uterus and placenta shifted markedly across gestation at E9, E15, and E18 (n = 3-8 dams that contributed uterus, cervix and placenta; one-way ANOVA with Tukey’s multiple comparisons; mean ± SEM, \**p<0.05*). Each data point in b and c represents the difference in PER2 peak time of the cervix and uterus or placenta explanted from the same dam. All surgeries were between ZT 2-6.

### Placental layers show intrinsic and coordinated circadian rhythms

To determine whether maternal or fetal layers contribute to circadian rhythms in the placenta, we explanted the placenta and compared PER2 bioluminescence in the intact placenta and isolated placental layers. We found that the maternal decidua, fetal junctional zone, and fetal labyrinth zone each exhibited circadian rhythms when isolated from each other at E15 or 18 with the decidua reliably peaking in the early subjective night (Supplementary Figure S9). Rhythms in the junctional and labyrinth zones were lower amplitude and less reliable in their times of peak expression. The isolated whole placenta exhibited circadian rhythms at all gestational ages examined (Supplementary Figure S9). To ensure viability of E18 cultured tissues, we added dexamethasone on the fifth recording day and found that all placental explants increased PER2 levels and circadian amplitude (Supplementary Figure S10). These data indicate that fetal and maternal layers of the placenta change their circadian cycling in response to exogenous glucocorticoid input and implicate the high amplitude circadian rhythms of the maternal layer in daily coordination of the placenta.

To evaluate the potential for maternal-fetal communication within the placenta, we chose to examine placenta explanted on E12 when fetal and maternal layers have matured. We found PER2::LUC rhythms were bright enough to be imaged with an ultra-cooled camera and the E12 placenta showed a daily wave of PER2 bioluminescence from maternal to fetal layers with higher amplitude circadian rhythms in the maternal placenta (Figure 5a). To directly test for circadian communication between layers, we treated E12 whole placenta with vehicle (0.01% ethanol), a glucocorticoid agonist (100 nM dexamethasone), or a progesterone/glucocorticoid antagonist (1 µM mifepristone). We found that Mifepristone decreased synchrony among pixels within each placenta compared to vehicle or dexamethasone (Figure 5b, c, *p* < 0.05; Mif, n = 5 explants, 530 ± 234 pixels/explant; Veh, n = 4, 370 ± 100 pixels/explant; Dex, n = 5, 314 ± 190 pixels/explant). These data, consistent with PER2::LUC expression recorded with photomultiplier tubes from the intact placenta and isolated placental layers, indicate that the placenta responds directly to glucocorticoid stimulation with synchronized circadian rhythms and that the decidua may coordinate circadian timing across the placenta.

**Figure 5.**
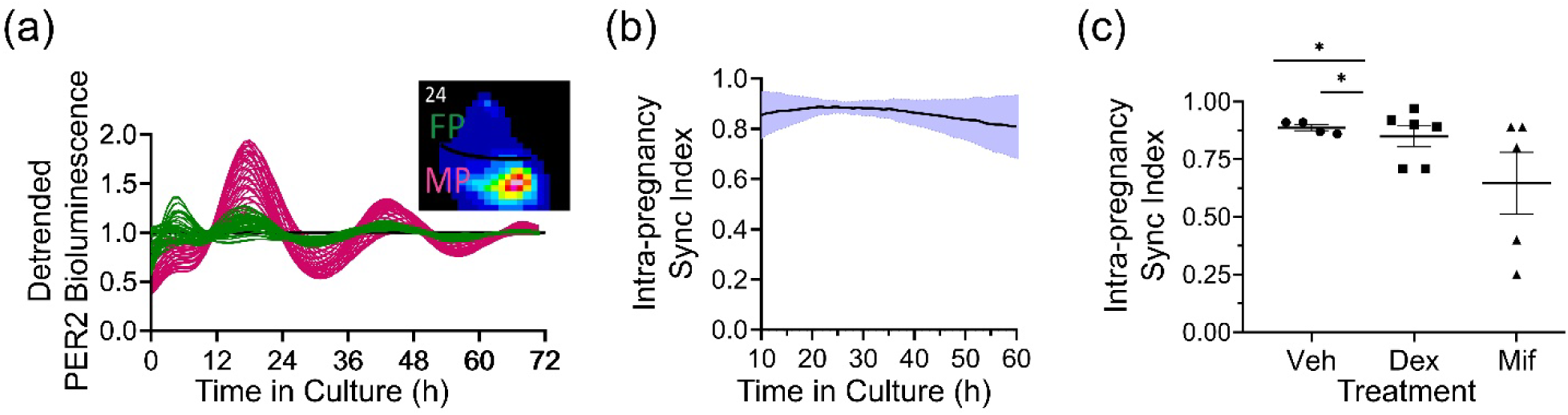
Glucocorticoid signaling synchronized circadian rhythms between the maternal and fetoplacental tissue *ex vivo*. (a) PER2 bioluminescence recorded over 3 days from 50 representative pixels within maternal (red) and fetal (green) E12 placenta. Note the high-amplitude daily PER2 rhythms in the maternal placenta that peaked at similar times as fetal placenta. Inset: Representative PER2::LUC bioluminescence image of the maternal (MP) and fetal (FP) placental layers after 24 h in culture. (b) Intra-pregnancy sync index indicated stable circadian synchrony between maternal and fetal placenta (mean ± SD, n = 4 explants). We computed the sync index (Kuramoto order parameter) from the wavelet-based phase estimation of PER2 signals across the tissue (370 ± 100 pixels/explant, mean ± SD). (c) The glucocorticoid and progesterone receptor antagonist, Mifepristone (Mif, n = 5 explants, 530 ± 234 pixels/explant) reduced synchrony across the fetoplacental pixels compared to vehicle (Veh, n = 4, 370 ± 100 pixels/explant) and dexamethasone (Dex, n = 5, 314 ± 190 pixels/explant, one-way ANOVA with Tukey’s multiple comparisons, mean ± SEM, \**p<0.05*).

We conclude that glucocorticoid signaling modulates synchrony between the maternal and fetal layers of the placenta.

## Discussion

The ontogeny of circadian rhythms has been difficult to resolve. By developing techniques to record *in utero* PER2 bioluminescence restricted to the conceptus, we find evidence indicating that fetoplacental PER2 expression increased during gestation with circadian modulation that gradually synchronized to peak around dusk by E15.5 in freely moving and in anesthetized dams. Glucocorticoid administration *in vivo* enhanced synchrony in daily rhythms of the maternal and fetal tissues which can be disrupted *in vitro* by glucocorticoid antagonists. We found that providing luciferin in drinking water dramatically avoided pregnancy complications including loss of *in utero* circadian rhythms and failure to deliver. We detected fetoplacental PER2 bioluminescence when we started recordings typically on E8.5. We conclude that the sensitivity and temporal resolution offered by recording a bioluminescent reporter of clock gene expression enabled the detection of circadian synchronization which has been difficult to evaluate by other methods.

Importantly, camera- and PMT-based methods reliably detected daily PER2 bioluminescence rhythms *in utero* by E15. These results indicate circadian PER2 expression in the fetus earlier than prior *in vivo* measures of fetal *Per2* mRNA, based on pooled data from multiple fetuses and pregnancies sacrificed at defined embryonic stages, which found no rhythms *in utero* or evidence for weak daily rhythms one or two days before birth (Dolatshad et al., 2010a; Houdek & Sumová, 2014; Kováciková et al., 2006; Sládek et al., 2004). Our findings are, however, consistent with prior *in vitro* research that found several fetal tissues expressed PER2 rhythmically as early as E13.5, but could not rule out that rhythms were induced by surgical isolation (Carmona-Alcocer et al., 2018; Dolatshad et al., 2010a; Landgraf, Achten, et al., 2015). Together with our real-time recordings of PER2::LUC expression from pregnant dams in constant darkness, we conclude that fetal-derived tissues are circadian *in utero*.

Across pregnancies, fetoplacental PER2 bioluminescence gradually synchronized to the local light cycle, peaking reliably around dusk by around E15.5. This timing corresponds with the rapid decline in 11 beta-Hydroxysteroid dehydrogenase type 2 (11β-HSD2) levels between E14.5 and E16.5 (Brown et al., 1996) reducing corticosterone deactivation and allowing high levels of maternal glucocorticoids (Wharfe et al., 2016) to reach the fetus (Burton & Waddell, 1999). Concordantly, we found that daily glucocorticoid injections to the dam, which can saturate 11β-HSD2 and interact with the fetus (Astiz et al., 2020a), accelerated circadian synchrony among fetoplacental units *in utero*. Of the 30 pregnancies monitored, 75% of all successful pregnancies exhibited circadian rhythms in PER2 bioluminescence and 83% of all failed pregnancies were also arrhythmic (Supplementary Tables S1, S2). This may indicate that failure of pups to become circadian and synchronize to their mom puts the pregnancy at risk. Interestingly, dexamethasone injections given out-of-phase with the normal rise in glucocorticoids in mice or humans has been associated with increased anxiety-like behaviors (Astiz et al., 2020a). We found that the response of fetoplacental PER2 rhythms depended on when the corticosterone was first injected relative to peak of PER2. This is consistent with phase-dependent effects of dexamethasone on shifting circadian rhythms in other tissues (Balsalobre et al., 2000; Spencer et al., 2018). We conclude that injections of glucocorticoids in late gestation can shift fetal circadian rhythms, and daily administration may entrain fetal circadian rhythms, making them potential candidates for maternal-fetal circadian communication. These data warrant studies to further test manipulations of maternal glucocorticoid secretion and the benefits of timed dexamethasone delivery. While adrenalectomy and glucocorticoid replacement can put a pregnancy at risk (White et al., 1980), timed optogenetic or pharmacological approaches (Astiz et al., 2020b; Astiz & Oster, 2020; Jones et al., 2021; Lehmann et al., 2023) to clamp corticosterone levels in mice could identify gestational windows when maternal corticosterone is necessary or sufficient to synchronize circadian rhythms in offspring and improve birth outcomes. Ultimately, our findings in mice suggest that delivering dexamethasone around the time of waking (when cortisol typically peaks) could improve maternal-fetal circadian synchrony, reduce the number of injections needed for healthy fetal lung maturation, and avoid the risk of complications including exogenous Cushing syndrome, weight gain, hypertension, and hyperglycemia (Fee et al., 2023; Ogueh & Johnson, 2000; Thevathasan & Said, 2020).

*In vitro*, waves of PER2 bioluminescence across the layers of the E12 placenta suggest maternal-fetal circadian coordination. With the caveats that our surgical isolation cannot perfectly segregate the placenta by layers and surgery could induce circadian rhythms that differ from *in vivo*, we found that all three placental layers have intrinsic PER2 rhythms as early as E15. Consistent with prior results which reported circadian rhythms intrinsic to the uterus and placenta during pregnancy (Akiyama et al., 2010; Čečmanová et al., 2019; Wharfe et al., 2011; Yaw et al., 2021), we found that the decidua had higher amplitude rhythms than the junctional zone and the labyrinth zone while the damped circadian rhythms of the fetal placenta increased in amplitude following dexamethasone stimulation. Finally, we found that daily rhythms in the uterus consistently peaked a few hours before the cervix and placenta *in vitro*, but around E15 these relationships became unreliable within and between pregnancies. Thus, our data, collected throughout gestation suggest a reorganization of the maternal and fetal circadian systems around E15 when we find increased maternal-fetal circadian synchronization. Nevertheless, our findings aligns with our results from *in utero* imaging indicating that daily rhythms across fetoplacental units become detectable between E15 and E16 depending on the pregnancy and the synchrony index within a pregnancy increases after E16. Prior studies

Our findings implicate glucocorticoids from the mother may synchronize daily rhythms in fetal tissues. A functioning maternal circadian system during gestation has been shown to be important in normal development including the initiation of daily rhythms in sleep (Landgraf et al., 2015; Mendez et al., 2012a, 2016; Summa et al., 2012a; Varcoe et al., 2016). Environmental factors (e.g., shift work, constant light, or repeated 6-h shifts in the light cycle) have been associated with disrupted maternal daily rhythms (e.g., in hormones or body temperature), placental abnormalities and failed pregnancies in rodents (Mendez et al., 2012b; Summa et al., 2012b) and increased risk of preterm birth, placental complications, and pediatric obesity in humans (Abeysena et al., 2009; Cai et al., 2019; Kader et al., 2021; Kawai, 2022; Knutsson, 2003; Zhu et al., 2004). We found highly synchronized daily rhythms between maternal and fetal layers of the E12 placenta that were sensitive to glucocorticoid agonist or antagonists *in vitro*, consistent with a prior observation that glucocorticoids induce and shift E17 maternal and fetal placental circadian rhythms depending on the time of DEX administration (Čečmanová et al., 2019). This suggests glucocorticoids synchronize daily rhythms within the placenta at least at E12 and E17. However, mifepristone also blocks progesterone signaling so we cannot rule out that progesterone or other signals may also serve to synchronize placental daily rhythms (Zhang et al., 2006). Future evaluation of the necessity and sufficiency of glucocorticoid signaling in maternal-fetal circadian synchrony should be placed in the context of other candidate maternal-to-fetal signals (Bates & Herzog, 2020). For example, single injections of melatonin or a dopamine receptor agonist on E15 can shift the subsequent locomotor rhythms of hamster offspring, suggesting they may suffice for *in utero* entrainment of fetal rhythms (Viswanathan & Davis, 1997) and extirpations of the maternal pituitary, adrenals, thyroid, ovaries, or pineal between E0-9 did not alter day-night glucose uptake measured in rat fetal SCN on E19 (Reppert & Schwartz, 1986) suggesting that multiple hormones or body temperature could suffice for maternal-fetal synchrony. Further studies should provide mechanistic insights into the cells and signals mediating synchronization of maternal and fetal daily rhythms during pregnancy.

There are additional limitations to the interpretation of results from *in utero* monitoring methods reported here. The exponential increase in PER2::LUC expression over gestation could arise with more fetal cells contributing to the PER2 signal or with augmented PER2 expression per fetal cell. Although cellular differentiation has been associated with released repression of key clock genes (e.g., increased expression of Per2 in pluripotent stem cells) in early-stage embryos (Dolatshad et al., 2010b; Inada et al., 2014; Umemura et al., 2014, 2017b), future studies should resolve how PER2 expression increases during gestation. Although we found preliminary evidence for day-night differences in fetal-derived PER2 expression on E8.5, we did not attempt recordings with higher temporal resolution in early gestation. Because most murine cells have either differentiated or fate-committed by that stage, future *in utero* monitoring of clock genes and proteins with cell-type specific reporters as cells proliferate and differentiate should advance our understanding of when, where, and how cells become circadian in the fetus. Because of fetal movement and growth, we could not distinguish the contributions of individual pups or tissues to the emergence of daily PER2 rhythms or their synchronization with each other and the mother. For example, much of the PER2 bioluminescence in early gestation derives from the fetal liver and the developing placenta.

We conclude that fetoplacental circadian rhythms progressively mature and align with maternal- and light-driven cues, likely through mechanisms involving glucocorticoids. Maternal circadian disruption during pregnancy, such as that caused by shift work, could impair maternal-fetal circadian synchrony and reproductive outcomes.

## Materials and Methods

### Animals

All experiments used mice maintained on a hybrid C57BL/6JN background. Animals were housed in a 12h:12h light-dark cycle (lights on from 7 AM-7 PM, Zeitgeber Times, ZT 0-12) in the Danforth Animal Facility at Washington University in St. Louis. Animals received *ad libitum* access to food and water throughout the experiments. Timed pregnancies were generated by pairing one male with two females overnight. Vaginal plugs observed the following morning defined embryonic day 0.5 (E0.5) and E1 ended with following lights off. All procedures were approved by the Animal Care and Use Committee of Washington University and followed National Institutes of Health guidelines.

### Imaging and analysis of *in utero* PER2::LUC bioluminescence in anaesthetized mice

To image fetoplacental PER2 *in utero*, we mated wildtype C57BL/6JN females to homozygous PER2::LUC transgenic males so that PER2::LUC is expressed only in the fetoplacental tissues of the pregnant wildtype dam. Luciferin was administered to pregnant dams either by intraperitoneal injection (10 mM D-luciferin, Gold Biotechnology, 10 min before imaging) or through the drinking water (10 mM D-luciferin, Gold Biotechnology) starting 24 h before imaging (Xenolight, PerkinElmer). We replaced luciferin-containing drinking water every 3 days. We removed abdominal area fur of the dam with depilatory cream (Nair, Church & Dwight) 48 h before recording. We anesthetized dams with 2% isoflurane in 1 L/min O₂ and imaged (IVIS Lumina III system, PerkinElmer) their ventral side 10 minutes after luciferin injections (where applicable). We used the same camera settings (15 cm field of view, 8x8 binning, f-stop 1, 10-min exposure from E8.5-11.5 and 2-min exposure from E12.5-E17.5) to image each dam twice daily from E8.5-E14.5 and every 4 h from E14.5-E17.5.

Bioluminescence was quantified from the captured images as total photon flux (photons/s) using fixed-area regions of interest (ROIs) drawn over the abdomen in Living Image 2.6 software (PerkinElmer).

### Continuous recording and analysis of *in utero* PER2::LUC bioluminescence in freely moving mice

To compare fetoplacental PER2 expression in mice that were not handled or anesthetized, wild-type dams carrying PER2::LUC conceptuses were generated as described above. Pregnant dams were provided drinking water containing 2.18 mM CycLuc1 starting 24 h prior to recording and housed individually in constant darkness from E13.5 in a light-tight box (Lumicycle In Vivo, Actimetrics) equipped with two photomultiplier tubes (Hamamatsu H8259-01) (Martin-Burgos et al., 2022). Drinking water was refreshed every 3 days. Fur on the abdominal area was removed with depilatory cream (Nair, Church & Dwight) 48 h before recording. In another set of experiments testing whether drinking induced daily rhythms in bioluminescence, a non-pregnant homozygous PER2::LUC (P2L/P2L) female was recorded under the same conditions with 10 mM D-luciferin supplemented in drinking water either *ad lib* or during the day. Dark counts were collected for 1 min every 15 min by closing the programmable shutter. Bioluminescence counts per minute from the two sensors were summed after subtraction of dark counts.

### Circadian analysis of *in utero* PER2::LUC bioluminescence

To test for daily cycling of *in utero* bioluminescence, we calculated a 24-h running mean of the bioluminescence and subtracted it from the raw signal. The resulting detrended traces were analyzed for circadian rhythmicity using MetaCycle (Wu et al., 2016), which integrates three algorithms (ARSER, JTK, and Lomb–Scargle). We defined traces as circadian if their period was between 18-30 h with a *p* < 0.05 value. Daily peak times and amplitude (peak-trough bioluminescence counts) were quantified using custom Python and MATLAB scripts (<u>GitHub -</u> <u>Herzog-Lab/Nikhil-Bates_JBR_2025</u>). We computed the inter-pregnancy sync index as the Rayleigh statistic R (Jammalamadaka & Sengupta, 2001), based on the daily bioluminescence peak times of pregnant dams. Rayleigh R ranges from 0 when rhythms peak at random times across animals to 1 when animals peaked at the same time of the day (Herzog et al., 2015). To assess whether fetoplacental units within a dam exhibit synchronous PER2, we computed intra-pregnancy sync index based on the PER2 bioluminescence signals from individual pixels from spatially aligned bioluminescence images collected every 4 h from E14.5–E17.5 using ImageJ (NIH). Briefly, we drew a fixed region of interest (ROI) around the abdomen, background subtracted it and, using a custom Python script (Granados-Fuentes et al., 2024), measured pixel intensities by implementing a multidimensional Gaussian filter in SciPy tools to generate time series of pixel bioluminescence. We detrended and smoothed the time series using Sinc filter and calculated the instantaneous phase using continuous wavelet transformation in pyBOAT (Mönke et al., 2020). Finally, we computed the intra-pregnancy sync index as the first-order Kuramoto order parameter by vectorial averaging of instantaneous phases on the complex plane (Mönke et al., 2020; Schmal et al., 2022). Kuramoto order parameter, like sync index ranges from 0-1 indicating complete desynchrony and total synchrony respectively.

### *In vivo* corticosterone treatment

Pregnant dams received daily subcutaneous injections of corticosterone (20 mg/mL; 50 mg/kg, Sigma-Aldrich) or vehicle (polyethylene glycol 400, Sigma-Aldrich) at ZT0 from E13.5–E17.5.

### *In vitro* PER2::LUC bioluminescence recording and analysis

In a separate cohort of PER2:LUC homozygous females, we recorded PER2 driven bioluminescence from maternal reproductive tissues for at least five days immediately after explantation at different gestational ages. Briefly, we sacrificed pregnant and non-pregnant females between ZT2-6 using CO₂ asphyxiation followed by cervical dislocation at E9, E12, E15, E18 and during metestrus and proestrus, respectively. In non-pregnant females, we determined estrous stage by collecting vaginal smears 2 h before tissue harvest and identifying cell types under bright-field microscopy. From each female, we isolated the placenta (where applicable), uterus, and cervix into chilled Hank’s buffered salt solution (HBSS, Sigma-Aldrich). Placentae were sliced longitudinally and either cut into cross-sectional explants containing all layers (maternal decidua, maternal–fetal junctional zone, and fetal labyrinth zone) or further dissected into individual layers (∼4 mm³). Uterine and cervical samples were cleared of fat and cut longitudinally into ∼2 × 2 mm pieces. All explants were placed on 0.4 μm MilliCell membrane inserts (Millipore) in 35-mm Petri dishes (BD Biosciences) containing 1 ml DMEM (Sigma-Aldrich, pH 7.2) supplemented with 25 U/ml penicillin, 25 mg/ml streptomycin (Invitrogen), 10 mM HEPES (Sigma-Aldrich), 2% B27 (Invitrogen), 0.35 g/L NaHCO₃ (Sigma-Aldrich), and 0.1 mM beetle luciferin (BioSynth). Dishes were sealed with vacuum grease (Yamazaki & Takahashi, 2005) and transferred to light-tight incubators maintained at 36°C and positioned under photomultiplier tubes (HC135-11MOD; Hamamatsu). We recorded bioluminescence from each explant in 10-min bins for 5 days. Bioluminescence traces with a period of 18–30 h and a correlation coefficient >0.70 were classified as circadian based on Chronostar (Maier et al., 2021) fits to days 2–5. Peak-to-trough amplitude was calculated from 36–60 h after recording onset using a custom MATLAB script (<u>GitHub -</u> <u>Herzog-Lab/Nikhil-Bates_JBR_2025</u>). We calculated the peak phase from the cosine fit (Chronostar) to days 2–5 and expressed in Zeitgeber Time (ZT; ZT0 = light onset time on the day of surgery). To test whether PER2 peaked at similar times of day across tissues or pregnancies, we applied the Rayleigh uniformity test where *p* < 0.05 indicates significant clustering (consistent timing) across tissues using Oriana software (version 4.02).

To analyze whether glucocorticoids affected circadian rhythms in the placenta, a subset of E12 placental cross-sections were placed in a light-tight incubator at 36°C under a CCD camera (Onyx, Stanford Photonics). These placentas were immediately cultured with either dexamethasone-a synthetic glucocorticoid (2 µL of 50 µM dexamethasone in 1 mL of recording media, 100 nM, Sigma-Aldrich), mifepristone–a glucocorticoid and progesterone receptor antagonist (1 µL of 1 mM mifepristone dissolved in ethanol, 1 µM, Sigma-Aldrich), or vehicle (0.01% ethanol in ddH2O). Images were collected every 30 minutes for 5 days and 2 images were averaged using ImageJ software (NIH) to generate one image per hour. Bright noise and cosmic radiation were filtered out using adjacent frame minimization in ImageJ and the movies were subjected to pixel-based analysis as described above.

### Additional statistical analyses

Sync indices were compared between corticosterone-injected and control mice using two-way ANOVA, while those across treatment groups were analyzed by one-way ANOVA followed by Tukey’s post-hoc comparisons. Unless specified, all statistical analyses and plotting were implemented using GraphPad Prism version (GraphPad Software, www.graphpad.com).

## Acknowledgements

The authors thank the members of the Herzog and England labs for valuable discussions and comments on the manuscript. This work was supported by National Institutes of Health Grants NINDS R01NS12116 and the March of Dimes Prematurity Research Center. KLN was supported by a fellowship from the McDonnell Center for Cellular and Molecular Neurobiology. Supporting Information

