## Supplemental Information for "Fetoplacental circadian rhythms develop and then synchronize to the mother *in utero*"

Short title: Maternal-fetal circadian synchrony

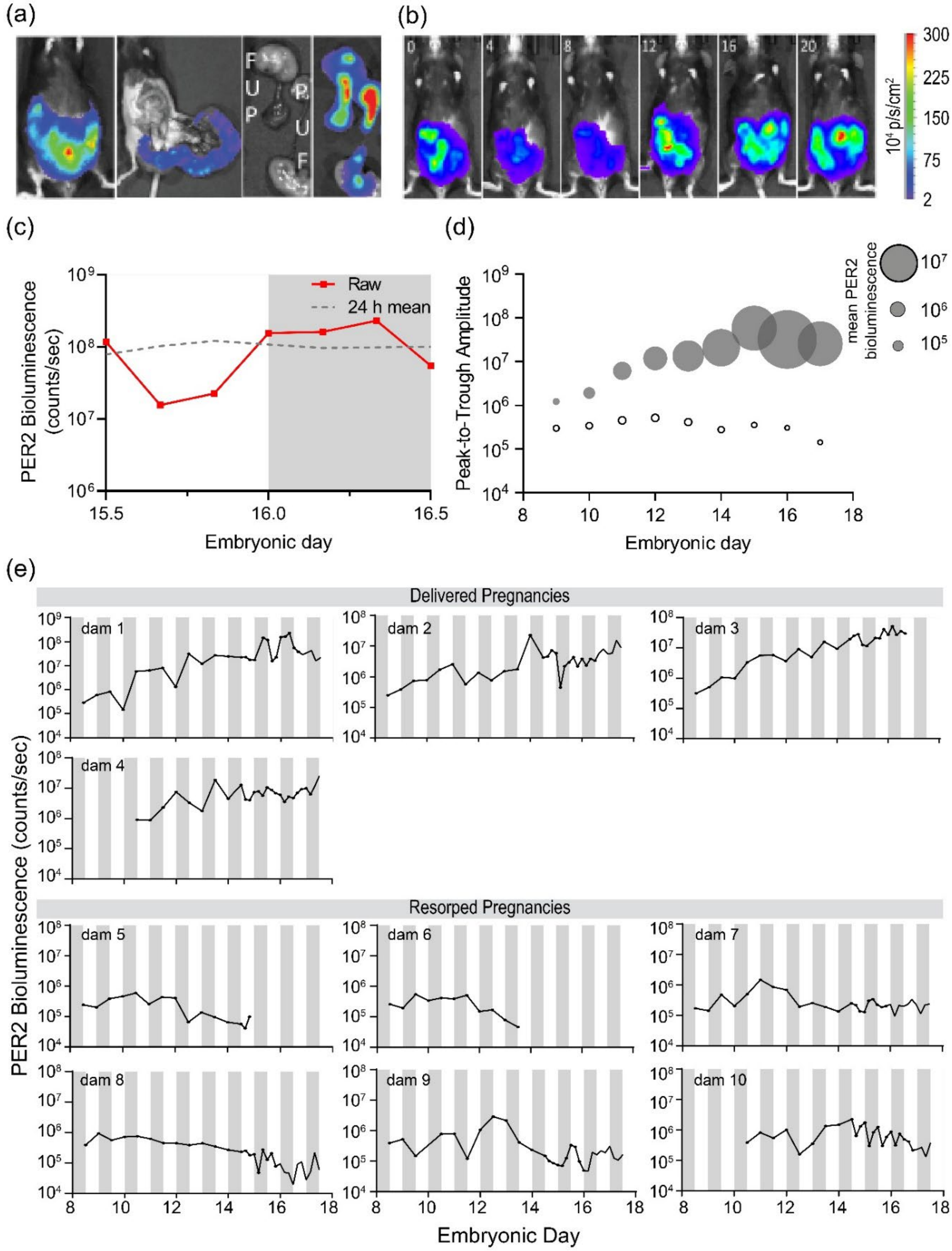


**Figure S1.** ***In utero*** **imaging of fetoplacental PER2 bioluminescence reveals that successful pregnancies are associated with gestational development of high-amplitude circadian rhythms.** (a) A representative dam showed fetoplacental PER2 bioluminescence 10 min after luciferin injection on embryonic day (E)17.5. After dissection, the bioluminescence localized to the two fetuses (F), their placentae (P), and umbilical cords (U). (b) PER2 bioluminescence imaged every 4 h (ZT0–20) on E17 in a representative dam varied with time of day. (c) PER2 bioluminescence (red) plotted around the 24-h running mean (dashed line) from the dam shown in (b). White and gray bars denote day and night, respectively. (d) Across gestation, daily peak-to-trough amplitude and mean (circle diameter) of fetoplacental PER2 increased in successful pregnancies (gray circles, n = 4). In failed pregnancies, amplitude and mean did not increase (open circles, n = 6). (e) Bioluminescence traces imaged following luciferin injections showed that fetoplacental PER2 increased through gestation in dams that delivered (n = 4, top) but not in pregnancies that failed (n = 6, bottom). White and gray bars denote day and night.

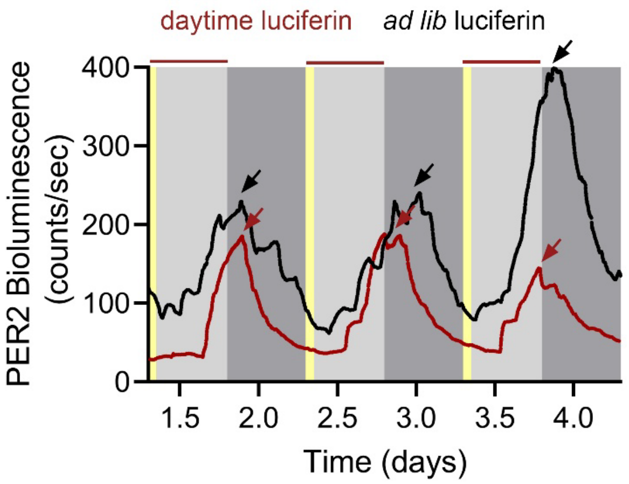


**Figure S2. Daily PER2 bioluminescence in a freely moving non-pregnant female mouse peaked at dusk and was minimally affected by the time of luciferin availability.** (a) PER2 bioluminescence recorded longitudinally in a non-pregnant, homozygous PER2::LUC female mouse that received 10mM D-luciferin-supplemented drinking water either *ad libitum* (black trace) or only during the day (red trace). The mouse was exposed to daily light between 7:00–7:15 AM (ZT0–0.25, yellow bands). PER2 bioluminescence was circadian under both conditions, with periods of 23.8 h (daytime luciferin) and 24.2 h (*ad libitum*), and peaked consistently around dusk (arrows).

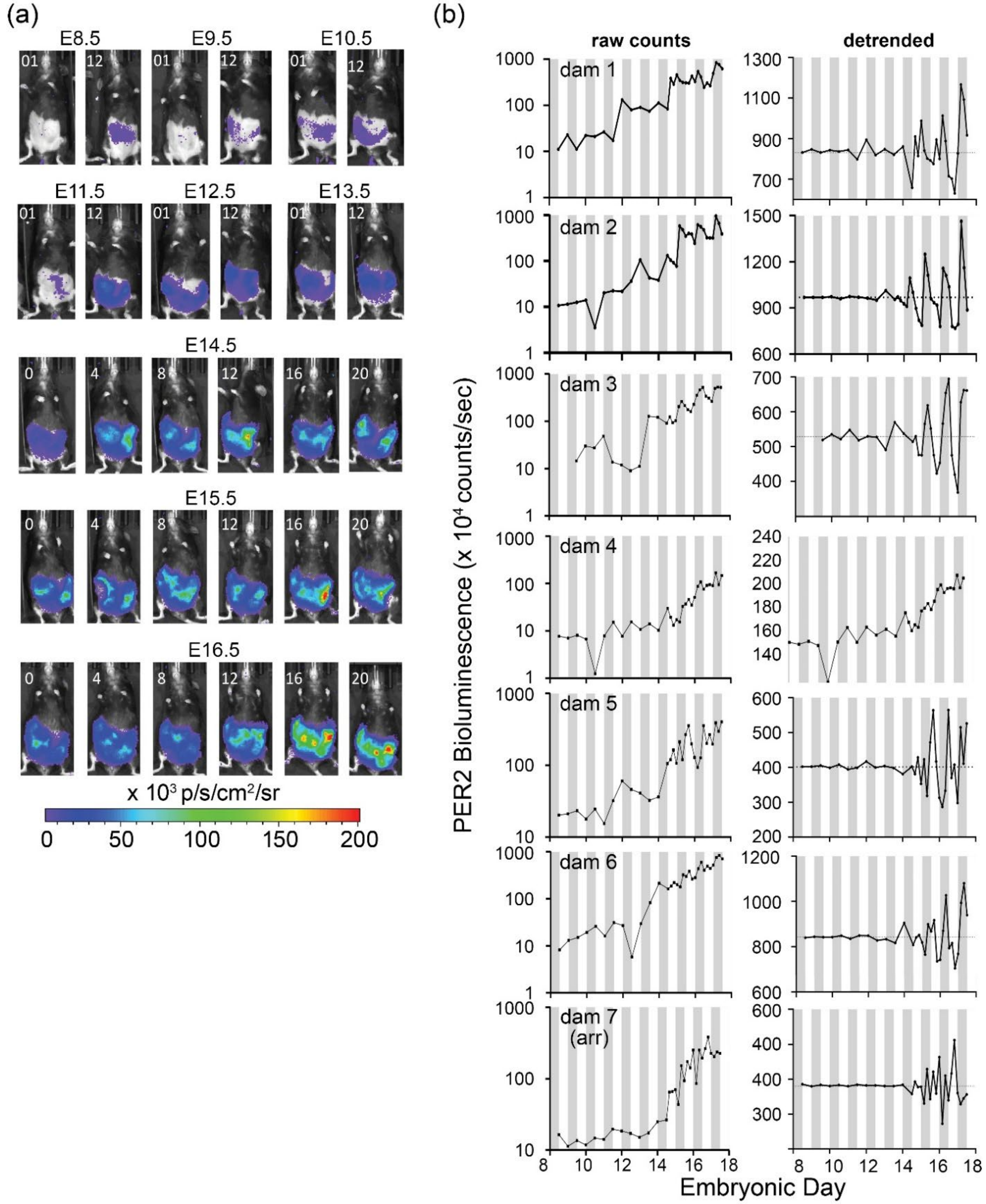


**Figure S3. Fetoplacental PER2 rhythms appear early, strengthen across gestation, and synchronize with maternal phase.** (a) Sequential PER2 bioluminescence imaged in a representative dam showed that fetoplacental PER2 was detectable at low amplitude on embryonic day (E)8.5, progressively increased throughout pregnancy, and varied across the day from dawn (Zeitgeber Time, ZT0; lights on) to dusk (ZT12; lights off). Dams received *ad lib* 10 mM D-luciferin-supplemented drinking water. (b) Raw counts (left) and detrended traces (right) of the fetoplacental PER2 bioluminescence imaged from all dams (n = 7). PER2 bioluminescence increased exponentially across gestation, with variable peak times until ~E15.5. Bioluminescence in 1 of 7 dams (arr) was not circadian (Metacycle, $p > 0.05$) and was excluded from analysis. For detrending, we subtracted the 24-h running average from each trace to reveal the first day when bioluminescence crossed the mean expression twice per cycle. High-amplitude rhythms and synchronous peak times emerged around E15.5 after which PER2 peaked around dusk (also see Figure S2). White and gray bars denote day and night.

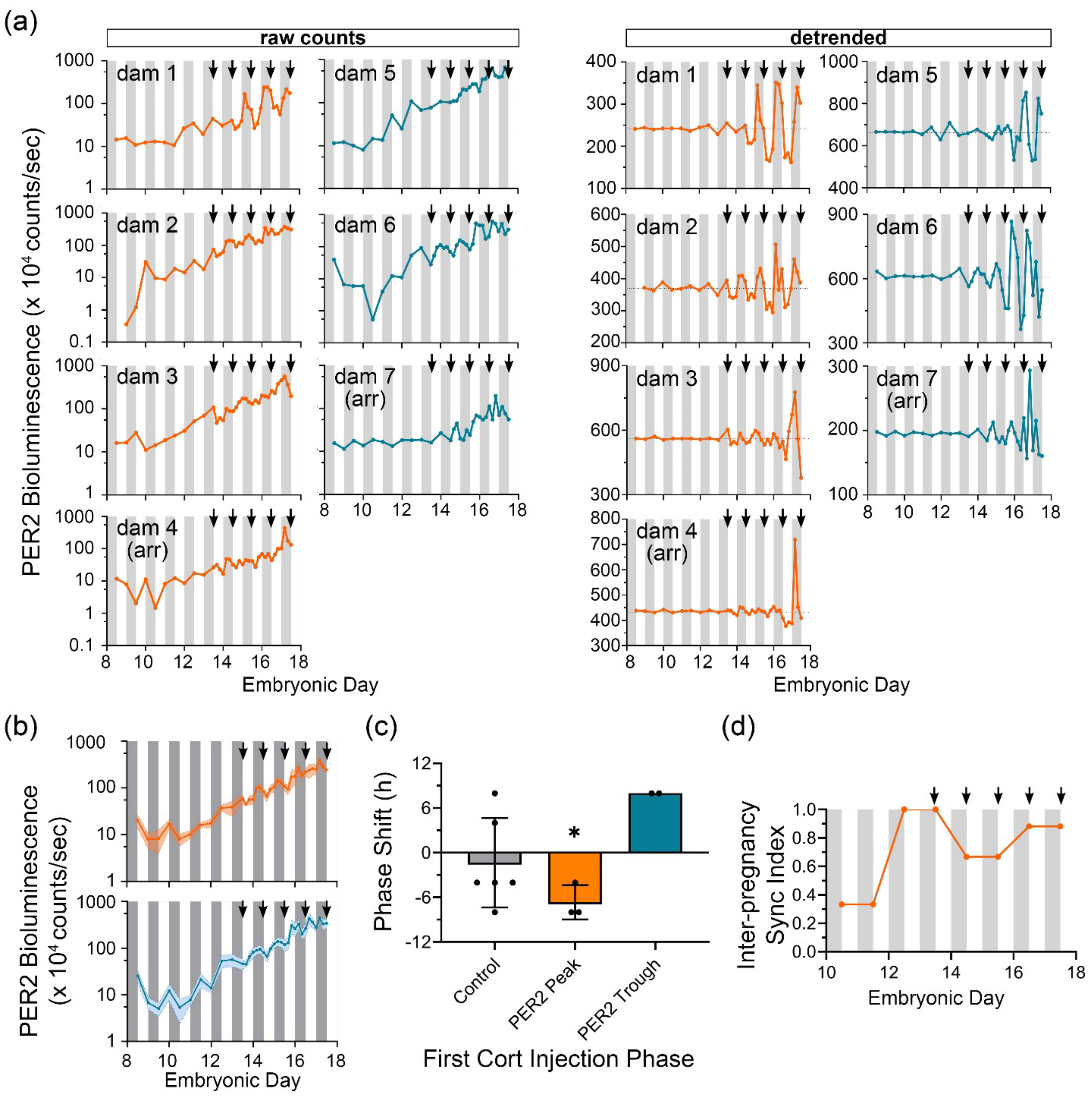


**Figure S4. Maternal corticosterone injections shift fetoplacental PER2 bioluminescence in a phase-dependent manner.** (a) Raw (left) and detrended counts (right) of PER2 bioluminescence imaged from E8.5 to E17.5 in 7 dams that drank D-luciferin-supplemented water and received daily corticosterone injections (arrows) at ZT0 starting on E13.5. Dams that received their first corticosterone injection near the peak (orange, n = 4) or trough (blue, n = 3) of their fetoplacental PER2::LUC expression all delivered. However, all but two showed circadian rhythms in fetal-derived PER2 bioluminescence (Metacycle, $p < 0.05$). (b) Of the dams that showed circadian rhythms *in utero*, average fetoplacental PER2 bioluminescence (mean ± SD) peaked during the day by E17 in dams that received their first

ZT0 corticosterone injection near the daily trough of PER2::LUC expression (blue, n = 2 dams) and at night in those initially treated near their PER2 peak (orange, n = 3 dams). Arrows mark corticosterone injections (E13.5–E17.5). (c) Corticosterone consistently delayed or advanced the peak of fetoplacental PER2::LUC expression from E13 to E17 by 4–8 h compared to the vehicle-treated controls ($*p < 0.05$, one-sample t-test, phase-shift significantly different from 0 h). (d) Daily corticosterone injections synchronized *in utero* PER2 bioluminescence peak times across pregnancies earlier than in controls (Figure S2). We calculated the inter-pregnancy sync index as the Rayleigh statistic of daily PER2 peak times across dams. White and gray bars denote day and night, respectively.

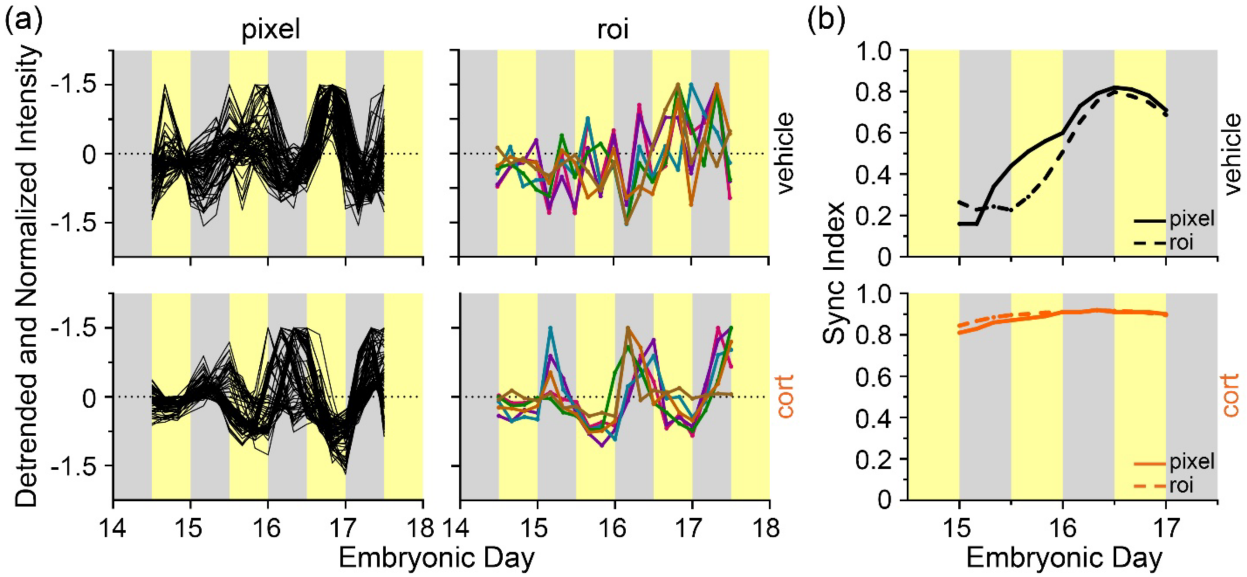


**Figure S5. Pixel-based and region-based analyses yielded similar measures of intra-pregnancy sync. (**a) Normalized and detrended fetoplacental PER2 bioluminescence imaged from E14.5 to 17.5 measured as pixel intensity (black line, left, n=100 representative pixels) from a representative control dam (top) and a corticosterone-injected dam (bottom). Bioluminescence from the same images of the same dams quantifed from 6 pup-sized regions of interest (ROI, 6 colored lines). (b) The intrapregnancy sync index among pixels (solid line) or pup-sized ROI (dashed line) were similar in both dams. Yellow and gray bars indicate day and night, respectively.

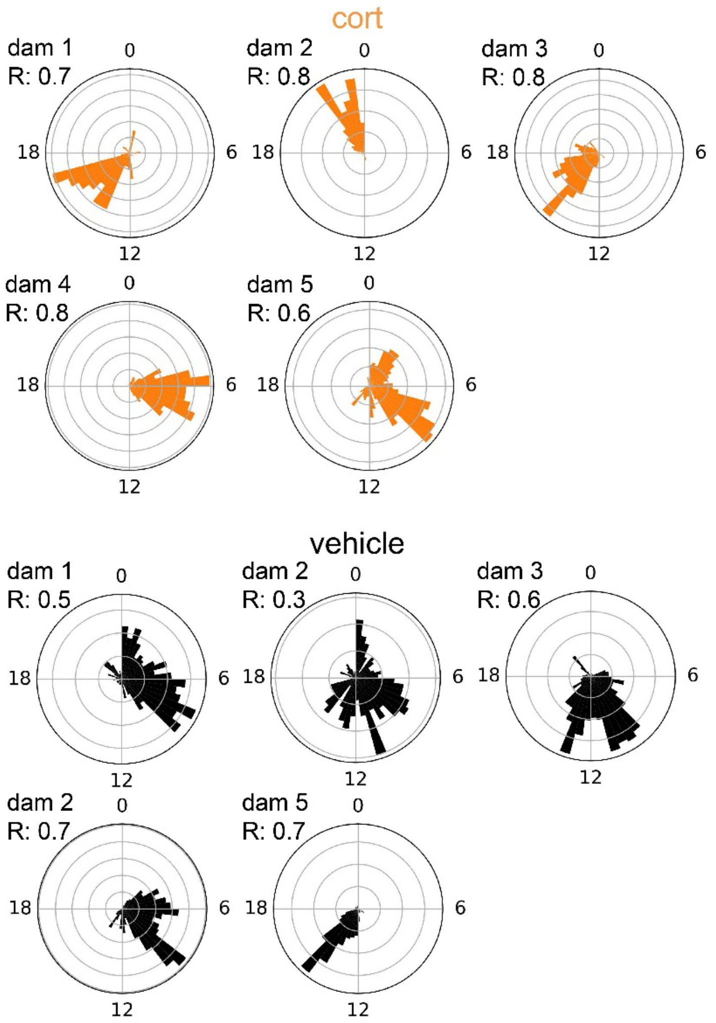


**Figure S6. Corticosterone synchronized PER2-driven bioluminescence across the fetoplacental tissue.** (a) Polar plots show time-of-day of the peak PER2 bioluminescence across pixels imaged on E15.5 from corticoterone- injected (orange, n = 5) and vehicle-injected dams (black, n = 5). The Rayleigh statistic (R) indicates the synchrony among pixels (0 = each pixel peaked at a random time of day; 1 = PER2 peaked at the same time of day in all pixels). Note that phases of pixel signals in corticosterone-injected dams had higher synchrony. Numbers around the circle indicate time of the day.

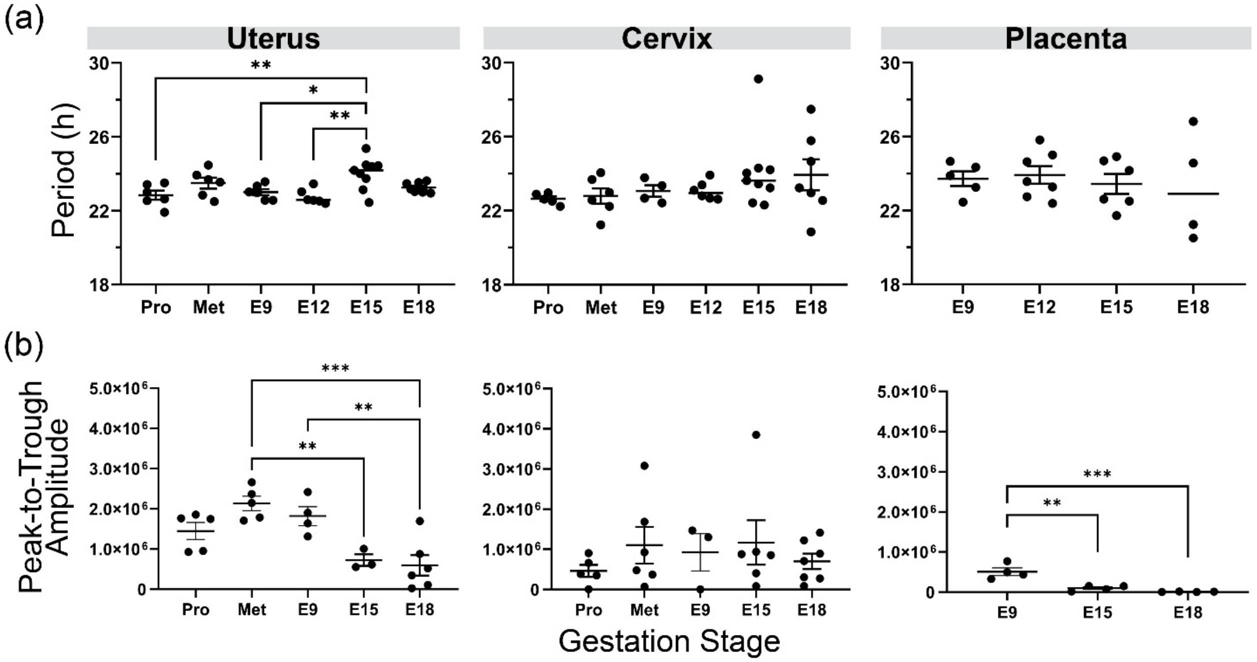


**Figure S7. Isolated maternal and placental tissues showed relatively stable circadian periods but decreasing amplitude with gestational age.** (a) Circadian period of PER2 bioluminescence did not change dramatically for the uterus, cervix, or placenta recorded at proestrus (Pro), metestrus (Met), E9, E12, or E18. At E15, the uterus had a longer circadian period. (b) The peak-to-trough amplitude of PER2 bioluminescence decreased in the last stages of pregnancy in the uterus and placenta explanted on E15 and E18 (One-way ANOVA with Tukey's multiple comparisons; mean ± S.E.M, *$p$<0.05, **$p$<0.01. **$p$<0.001).

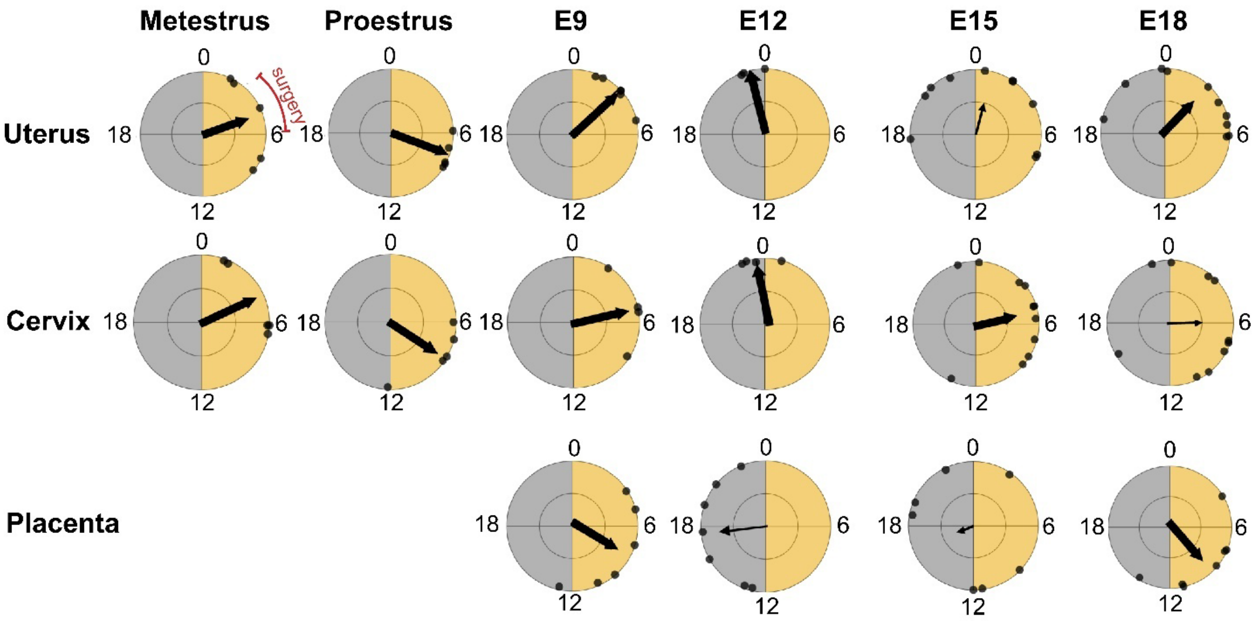


**Figure S8. Daily PER2 bioluminescence peak at reliable times across reproductive tissues depending on gestational age.** Rayleigh plots reveal the average time of day when PER2 luminescence peaked (arrow) in different tissues from 4-10 mice on the second day *in vitro*. Explants (black dots) from the uterus collected on metestrus, for example, expressed average peak PER2 around ZT4, 4 h after onset of light (yellow). Large arrows show samples that had times of peak expression that significantly differed from random (i.e., significant phase clustering, Rayleigh test). PER2 expression peaked at reliable times in the uterus at metestrus ($p = 0.04$), proestrus ($p = 0.0002$), E9 ($p < 0.001$), E12 ($p = 0.04$), and E18 ($p = 0.003$), but not E15 ($p=0.10$). PER2 expression in the cervix reliably peaked at metestrus ($p = 0.04$), proestrus ($p=0.02$), E9 ($p=0.04$), E12 ($p=0.01$), and E15 ($p=0.02$) but not E18 ($p=0.48$). In the placenta, PER2 expression reliably peaked at E9 ($p=0.047$) and E18 ($p=0.004$), but not E12 ($p=0.10$) and E15 ($p=0.87$). Yellow and gray regions indicate day and night. All explants were prepared between ZT2-6 (red).

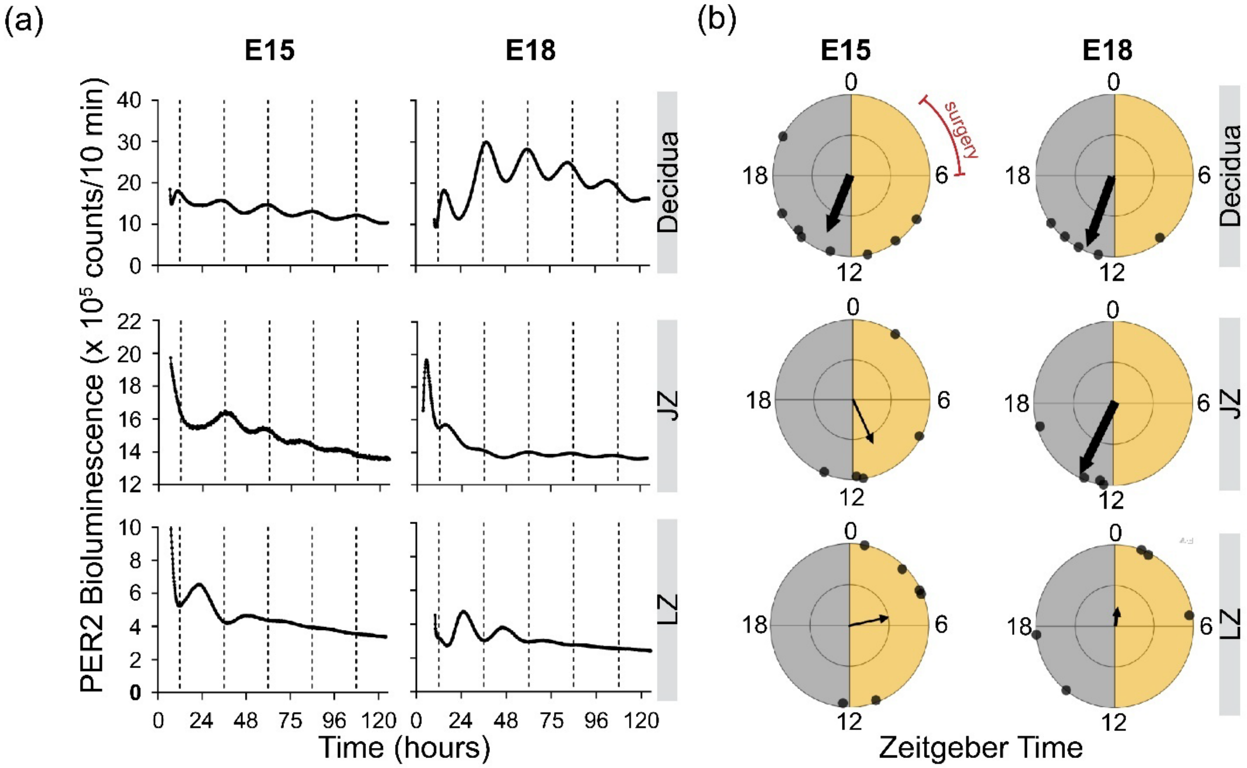


**Figure S9. Circadian rhythms in isolated placental layers.** (a) PER2 bioluminescence from representative explants of the decidua (top), junctional zone (JZ, middle), and labyrinth zone (LZ, bottom) collected at E15 or E18. All three layers exhibited intrinsic circadian rhythms. (b) Rayleigh plots show that PER2 in the decidua peaked, on average, in the early subjective night while the other two layers were less reliable in their time of peak PER2 bioluminescence. Yellow and gray regions indicate day and night. All explants were prepared between ZT2-6 (red).

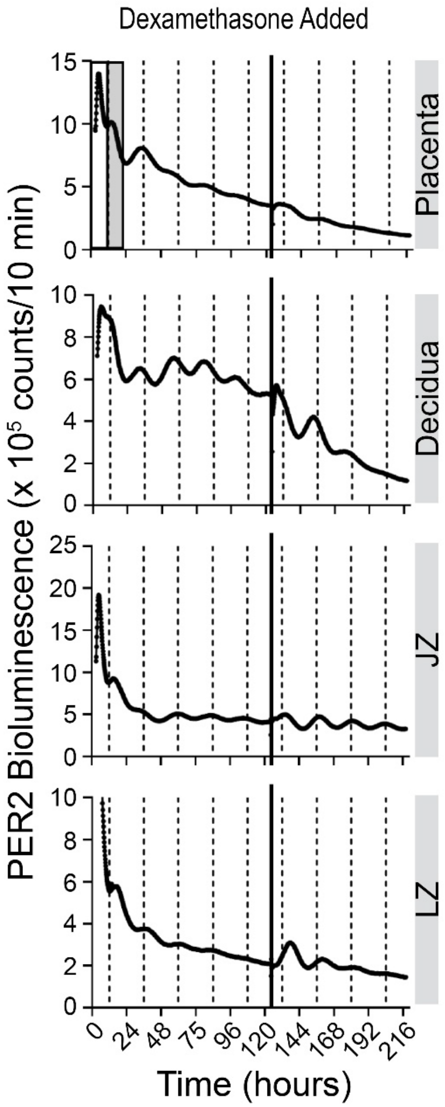


**Figure S10. Isolated placenta and placental layers remained viable after five days of ex vivo recording.** PER2 bioluminescence from representative cross sections of placenta, decidua, junctional zone (JZ), and labyrinth zone (LZ) collected at E18 show that all tissues responded to dexamethazone administration (solid vertical line) after 5 days in vitro with an increase in PER2 bioluminescence. Dashed lines mark every 12 h relative to the prior light-dark cycle (White-grey bar).

| Recording method | Luciferin Delivery | Fig. # | Total recorded dams | Circadian & Delivered | Arrhythmic & Delivered | Circadian & Resorbed | Arrhythmic & Resorbed |
|---|---|---|---|---|---|---|---|
| Continuous bioluminescence recording in freely moving dams | water | 1 | 6 | 6 | 0 | 0 | 0 |
| Imaging bioluminescence in anaesthetized dams | water | 2, S3, S4 | 14 | 11 | 3 | 0 | 0 |
| | injection | S1 | 10 | 1 | 3 | 1 | 5 |

**Supplementary Table S1. Summary of luciferin delivery methods and pregnancy outcomes.** Number of dams recorded under different luciferin delivery methods and their circadian and pregnancy outcomes.

| Pregnancy outcome | Fetoplacental PER2::LUC | Percent |
|---|---|---|
| Delivered | Circadian | 75.0 |

| | | |
|---|---|---|
| Resorbed | Arrhythmic | 83.3 |

**Supplementary Table S2. Summary of pregnancy outcomes and fetoplacental PER2 expression patterns across all experiments.**
